# Post-replicative lesion processing limits DNA damage-induced mutagenesis

**DOI:** 10.1101/2023.09.04.556208

**Authors:** Katarzyna H. Maslowska, Ronald P. Wong, Helle D. Ulrich, Vincent Pagès

**Author notes:** Present address: Institute of Molecular Biology (IMB), 55128 Mainz, Germany.

## Abstract

DNA lesions are a threat to genome stability. To cope with them during DNA replication, cells have evolved lesion bypass mechanisms: Translesion Synthesis (TLS), which allows the cell to insert a nucleotide directly opposite the lesion, with the risk of introducing a mutation, and error-free Damage Avoidance (DA), which uses homologous recombination to retrieve the genetic information from the sister chromatid.

In this study, we investigate the timing of lesion bypass and its implications for the accuracy of the process. Our findings reveal that DNA polymerase η can bypass common, UV-induced TT-cyclobutane pyrimidine dimers at the fork, immediately after encountering the blocking lesion. In contrast, TLS at TT(6-4) photoproducts and bulky G-AAF adducts, mediated by Rev1 and Pol ζ, takes place behind the fork, at post-replicative gaps that are generated downstream of the lesion after repriming. We show that in this latter situation, TLS competes with the DA pathway, thus reducing overall mutagenicity of damage bypass. Additionally, our study demonstrates that Exo1 nuclease influences the balance between TLS and DA by modulating the size of the post-replicative gaps.

## Introduction

The DNA of every organism is continually subject to damage from various exogenous and endogenous agents. The resulting lesions will frequently block the progression of replicative DNA polymerases, impeding the progression of the replication fork and thereby posing a threat to genome stability. Cells have evolved DNA damage bypass mechanisms (also named DNA damage tolerance) that allow them to cope with DNA lesions during replication. Translesion synthesis (TLS) is a mostly error-prone process involving specialized DNA polymerases that insert nucleotides directly opposite the lesion. Damage avoidance (DA) is an error-free process relying on homologous recombination (HR) to bypass the damaged site. Both the choice of TLS polymerase and the balance between error-prone TLS and error-free DA determine the level of mutagenesis during lesion bypass.

DNA damage bypass in eukaryotes is regulated by ubiquitylation of the sliding clamp, PCNA (1). The buildup of RPA-coated ssDNA downstream of a replication-blocking lesion recruits the Rad6/Rad18 complex that attaches a single ubiquitin to K164 of the resident PCNA (2). This modification initiates the TLS pathway by enabling recruitment of the specialized polymerases to the damaged site through their ubiquitin-binding domains (3, 4). Further extension of this ubiquitin to a polyubiquitin chain by the Ubc13/Mms2/Rad5 promotes the Damage Avoidance pathway (1, 5, 6).

Whether cells deal with DNA lesions at the replication fork or post-replicatively has been a long-standing debate. Initial studies by Rupp and Howard-Flanders (7) proposed that repriming could occur downstream of a lesion, leading to the generation of gaps behind the fork that were later filled in post-replicatively. However, this perspective changed with the discovery of TLS polymerases: at that time, the prevailing model suggested that TLS polymerases would transiently replace the replicative DNA polymerase at the fork, without the need for repriming or formation of gaps (8). This model was challenged when gaps were directly observed using electron microscopy in UV-irradiated *Saccharomyces cerevisiae* (9), and replication restart downstream of a lesion was observed *in vitro* in *Escherichia coli* (10). Following these observations, several studies demonstrated that *S. cerevisiae* deals with a range of DNA lesions in a post-replicative manner (11-14).

Edmunds *et al.* have shown in DT40 cells that bypass can occur both at the fork and post-replicatively, the choice being regulated by PCNA ubiquitylation and Rev1 (15). The identification of PRIMPOL in mammalian cells strongly supports the repriming model and, consequently, post-replicative lesion bypass (16, 17).

While repriming is now generally accepted, there remains some controversy regarding the events that occur at the replication fork. TLS could in principle occur both at the replication fork and at a post-replicative gap. Similarly, DA can occur by Homologous Recombination (HR) at a post-replicative gap, but strand exchange could also take place directly at the replication fork through the formation of regressed fork, also known as a “chicken-foot” structure. Such structure could facilitate the use of the sister chromatid as a template without the need of generating post-replicative gaps (18).

In this study, we reconcile both models by investigating the timing of lesion bypass in *S. cerevisiae*. We show that for TT-CPD (cyclobutane pyrimidine dimer) lesions, TLS by DNA polymerase η (Pol η) can occur at the fork, rapidly after the encounter with the blocking lesion. We also show that for lesions that are mainly bypassed by Rev1-Pol ζ, such as the TT(6-4) (thymine-thymine pyrimidine(6-4)pyrimidone photoproduct) or G-AAF (N-2-acetylaminofluorene at the C-8 position of a guanine residue), TLS occurs behind the fork, at post-replicative gaps that are generated downstream of the lesion via repriming. We found that in this latter situation, TLS activity is limited by competition with the DA pathway. Finally, we show that the nuclease Exo1, by extending the size of the post-replicative gaps, favors damage avoidance over TLS, thus further limiting the extent of damage-induced mutagenesis.

## Results

### The experimental system

To investigate the timing of lesion bypass, we examined the processing of several replication-blocking lesions and their partitioning between TLS and DA bypass. For this purpose, we used a recently developed system that allows us to introduce a single lesion at a precise genomic locus in *S. cerevisiae* and monitor its bypass by either TLS or DA (19). Briefly, a non-replicative plasmid containing the single lesion of interest is inserted at a specific locus within the yeast genome using the Cre recombinase and modified lox sites. The chromosomal integration site is located close to an early replication origin, resulting in an encounter of the replication fork with the lesion early in S phase. As the lesion is located within the *lacZ* reporter gene, bypass is monitored by classifying colonies by color (Supp. Figure 1). Total bypass events (DA or TLS) are plotted as a percentage of the numbers resulting from the integration of a non-damaged vector. Hence, percentages lower than 100% reflect a failure to bypass the lesion.

Using this system, we had previously monitored the bypass of a common UV-induced lesion, the TT(6-4) photoproduct, which is primarily processed by a combination of TLS polymerases ζ (Pol ζ) and Rev1 (19). We had shown for this lesion that TLS competes with DA (19). This was evident because inactivation of *UBC13*, encoding the E2 responsible for PCNA poly-ubiquitylation and thus initiation of DA, led to a decrease in the use of DA from ∼100% to 58% and a concomitant 10-fold increase in the level of TLS from 4% to 42%, without significantly affecting cell survival. Inactivation of *RAD51*, encoding the recombinase in charge of DA, produced a comparable decrease in DA and concomitant increase in TLS (19). Surprisingly, we did not observe this competitive relationship when using the TT-CPD (19), a lesion that is bypassed primarily by Pol η (encoded by *RAD30*) and only partially by Pol ζ-Rev1. We therefore asked whether competition between DA and TLS depended on the nature of the lesion or rather on the identity of the TLS polymerase(s) responsible for its bypass.

### DA competes with Pol ζ-Rev1, but not with Pol η-dependent TLS

To address this question, we first monitored the bypass of the G-AAF lesion, which, like the TT(6-4) lesion, is predominantly processed by Pol ζ-Rev1 (although a minor fraction of the bypass also involves Pol η) (20, 21). As shown in Figure 1A, inactivation of either *UBC13* or *RAD51* led to a decrease in DA and an increase in TLS, indicating a competition between the two pathways. In the *ubc13* and *rad51* strains, some DA persisted as we still observed a significant number of white colonies. These could have arisen from *RAD51*-independent template switching mechanisms that rely on *RAD52* (22). Overall, these results are similar to those previously obtained with the TT(6-4) lesion (19).

**Figure 1:**
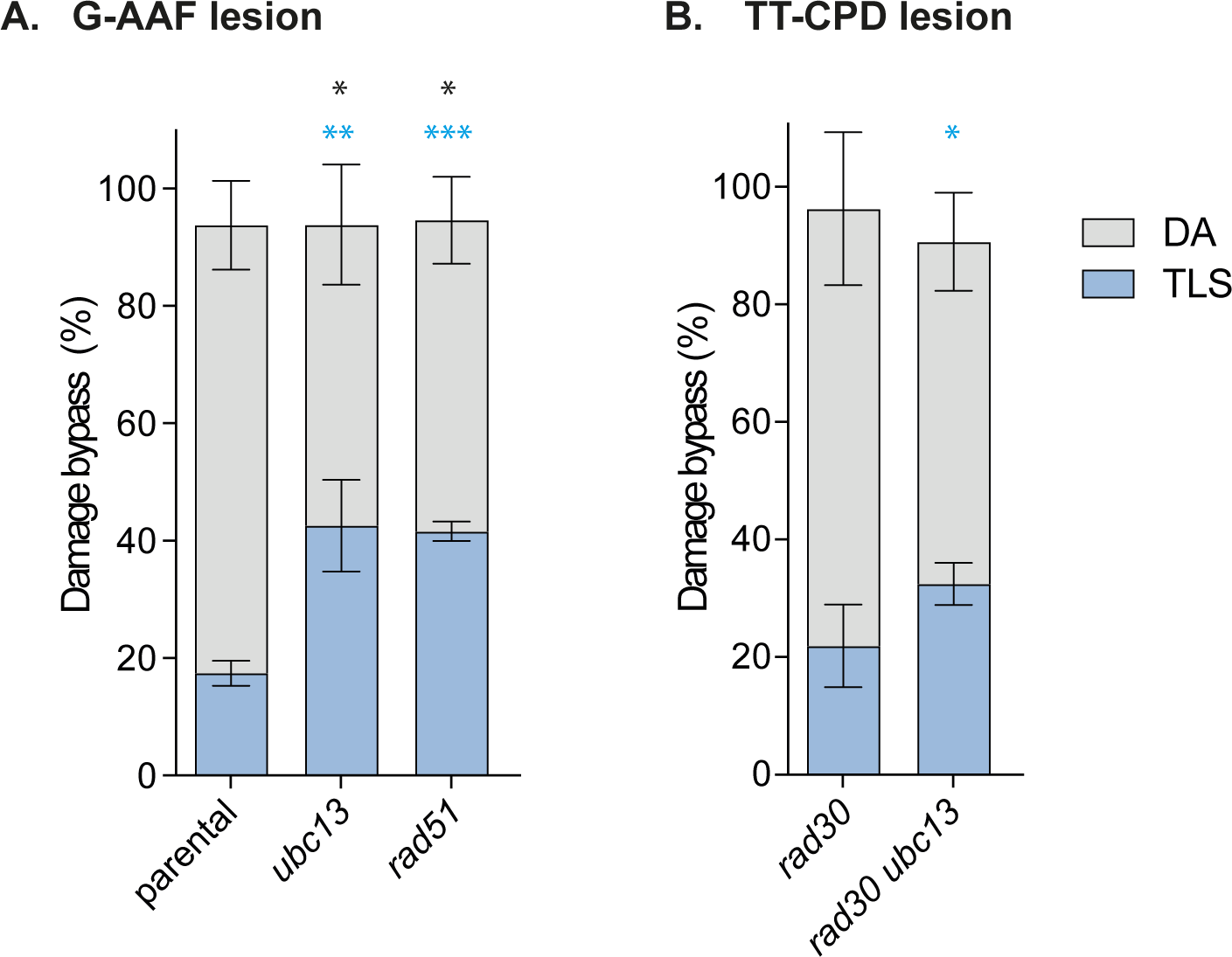
Partitioning of DNA damage bypass events. **A:** at a G-AAF lesion. **B:** at a TT-CPD lesion. Percentages represent lesion bypass compared to the non-damaged control. A bypass percentage below 100% reflects a lower survival upon integration of the damaged vector compared to the control. Unpaired t-test was performed to compare TLS and DA values from the different mutants to the parental strain (A) or to the *rad30* strain (B). (*p < 0.05; **p < 0.005; ***p < 0.0005).

We then re-assessed the bypass of the TT-CPD lesion in the absence of Pol η, i.e., under conditions where its bypass relies entirely on Pol ζ-Rev1 (Figure 1B). In this situation, we observed an increase in TLS when *UBC13* was inactivated, thus confirming a competitive relationship between DA and TLS when the damage is bypassed by Pol ζ-Rev1 rather than Pol η. We conclude that TLS competes with DA only when mediated by Pol ζ-Rev1, but not when Pol η mediates the bypass.

### Competition between TLS and DA is restricted to post-replicative bypass

Our results raised the question of why DA competes with Pol ζ-Rev1 activity, but not with Pol η-dependent bypass. Previous reports indicate that the expression level of Rev1 is significantly higher in G2/M compared to S phase (23, 24), while Pol η is expressed constantly throughout the cell cycle (24). At the same time, we had previously shown that the DA pathway mainly operates post-replicatively (13). We therefore hypothesized that Pol ζ-Rev1 might preferentially act at post-replicative gaps. Following the encounter with a replication-blocking lesion, a repriming event would then place Pol ζ-Rev1-dependent TLS in competition with DA. In contrast, bypass of TT-CPD lesions by Pol η might dominate directly at stalled replication forks without the need for repriming, thus avoiding the competition with DA.

If this model were valid, forcing Pol η to bypass the TT-CPD lesion post-replicatively rather than at the fork should induce a competition between TLS with DA for this lesion, as observed for the other two lesions tested.

To test this model by enforcing post-replicative action of Pol η, we restricted its expression to the G2/M phase of the cell cycle. We used two different strategies (Figure 2): i) we placed *RAD30* under control of the *CLB2* promoter and its proteasome-dependent cell cycle-regulated degron element (12), limiting its expression to the G2/M phase of the cell cycle (*clb2-RAD30*); ii) we used the auxin-inducible degron (AID*) system (25) to induce Pol η degradation in G1 and S phase, thereby restricting its expression to G2/M (*RAD30-AID\**).

**Figure 2:**
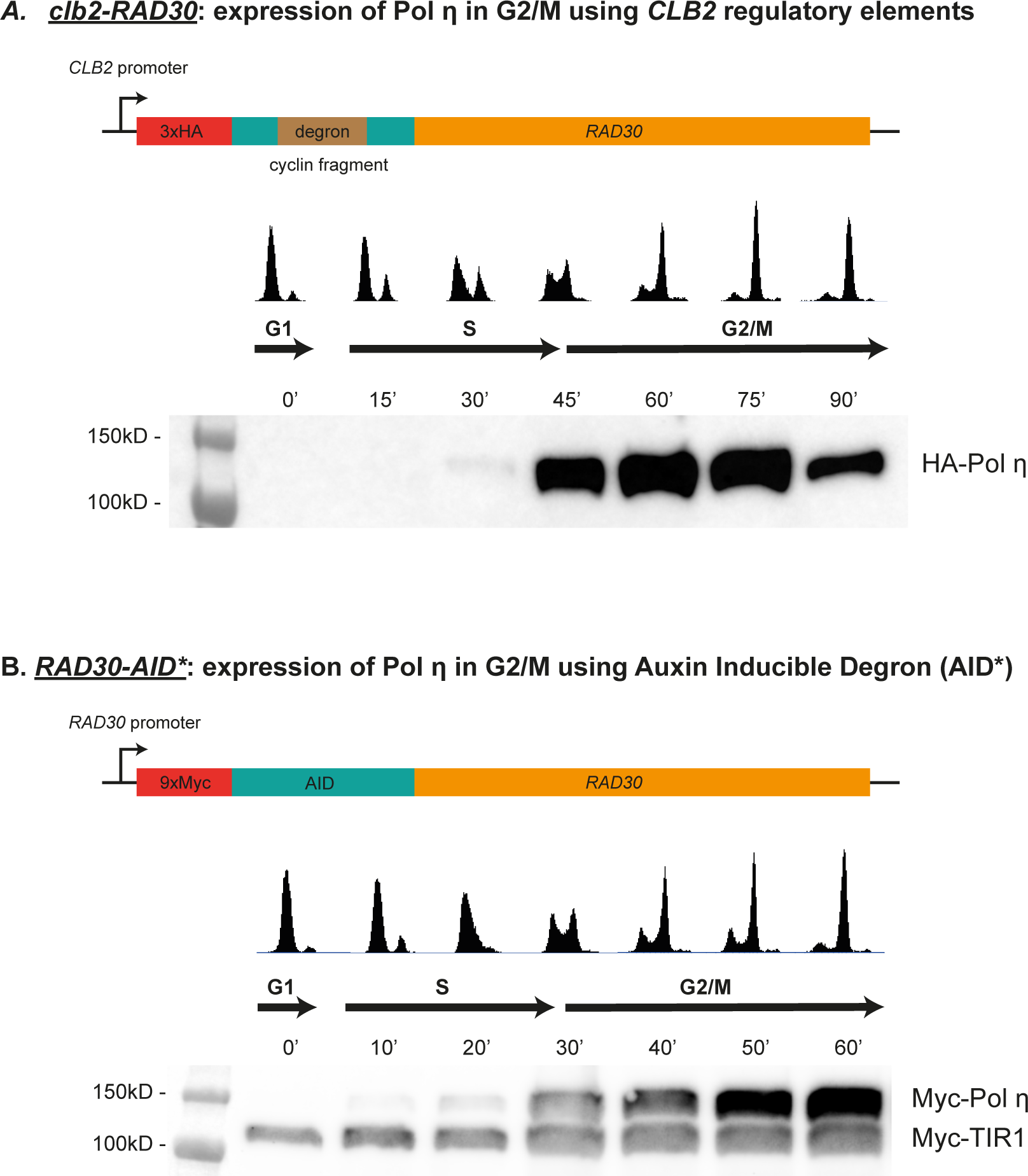
Constructs restricting Pol η expression to the G2/M cell cycle phase **A.** *clb2-RAD30*, allowing for regulation via the *CLB2* promoter and cyclin degron element. **B.** *RAD30-AID\**, allowing for regulation via the auxin-inducible degron. Auxin was present at 1 mM concentration during cell synchronization with alpha factor and was removed upon release of cells from the G1 arrest. Western blots show protein levels of HA-Pol η or Myc-Pol η at different phases of the cell cycle.

To verify the cell cycle regulation of Pol η, we synchronized cells in G1 using alpha-factor and monitored protein levels and cell cycle profiles upon release from the arrest by Western blotting and flow cytometry. As shown in Figure 2A, the *clb2-RAD30* construct afforded peak expression of Pol η in G2/M. However, while protein levels were low in G1 and early S phase, they already started increasing in mid-to-late S phase. *RAD30-AID\** (Figure 2B) showed a more effective restriction of expression during G1/S and a high expression level in G2/M. Another advantage of this construct is that it allows a physiological level of expression of Pol η as it is controlled by its native promoter.

Using these two strains, we measured the level of TLS and DA upon introduction of a single TT-CPD lesion into the genome during the S phase of the cell cycle. We accomplished this by electroporating the damaged vector immediately after washing away the alpha-factor and auxin. As shown in Figure 3A, compared to the parental strain, restriction of Pol η exclusively to G2/M led to a significant reduction in TLS at the TT-CPD lesion, accompanied by a simultaneous increase in the level of DA. It is important to note that the TLS level, although reduced, did not reach the same low levels observed in the complete absence of Pol η (Figure 1B). This could be due to a low level of Pol η already present in S phase that contributed to TLS. Alternatively, Pol η could continue to participate in TLS in G2/M even when in competition with DA, therefore resulting in an overall higher level of TLS than in the complete absence of the polymerase.

**Figure 3:**
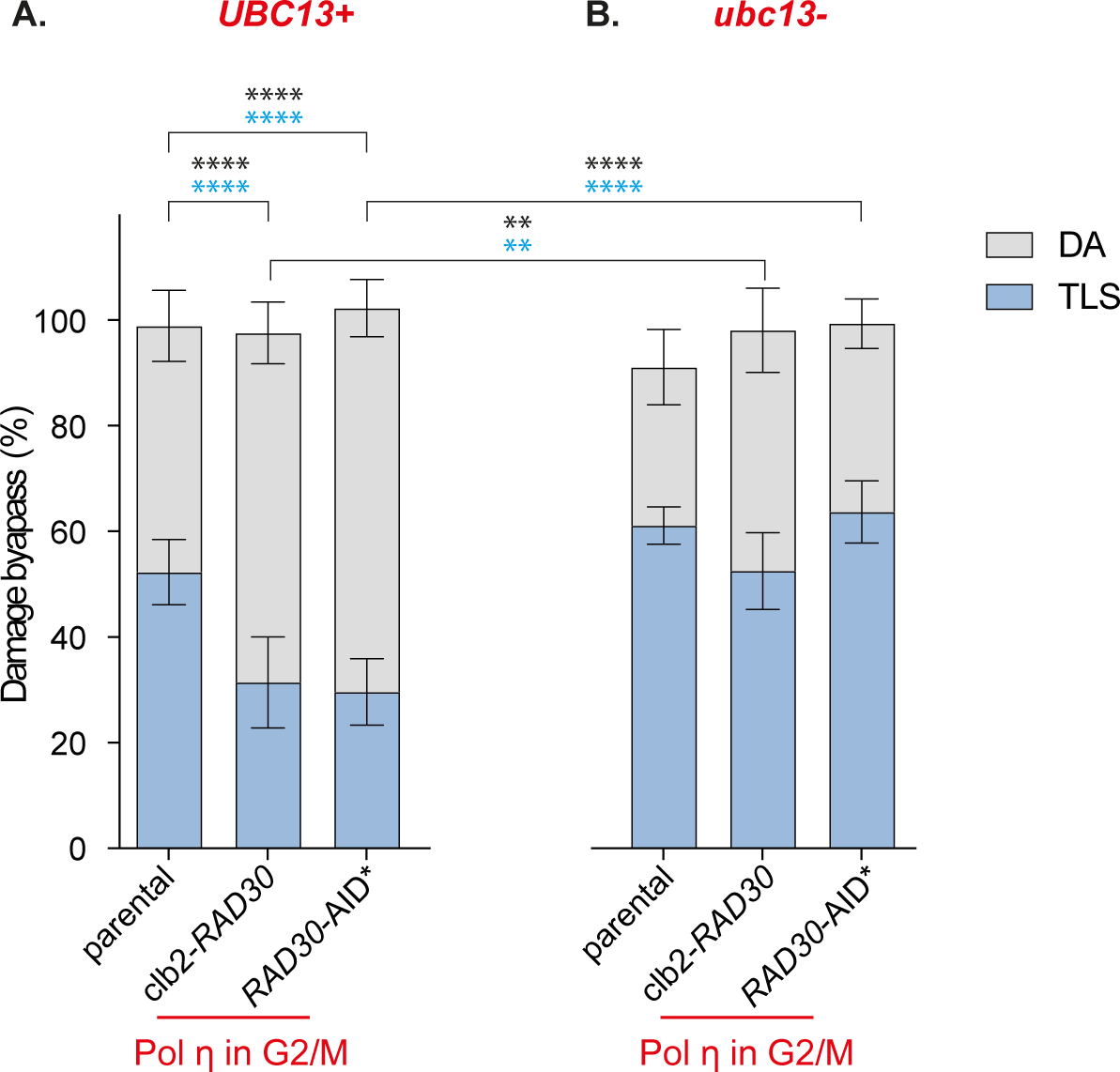
Partitioning of DNA damage bypass events at a single TT-CPD lesion in strains expressing Pol η only in G2/M. **A**. in cells proficient for DA (*UBC13+*), **B.** in cells deficient for *UBC13*-dependent DA (*ubc13*-). Unpaired t-test was performed to compare TLS and DA values from the different mutants to the parental strain. (*p < 0.05; **p < 0.005; ***p < 0.0005).

These observations supported our hypothesis that a post-replicative action of Pol η (in G2/M) would place it in competition with DA. To further validate this model, we repeated the experiment in a strain where *UBC13* was inactivated in order to prevent DA (Figure 3B). As previously shown (19), inactivation of *UBC13* in the parental strain where Pol η is expressed constitutively did not lead to a significant increase in TLS. However, when Pol η expression was restricted to G2/M, i.e., a situation where TLS potentially competes with DA, inactivation of *UBC13* caused an increase in TLS. Remarkably, in this situation, TLS was effectively restored to levels comparable to the parental strain with constant Pol η expression.

Taken together, these experiments validate our model. Thus, when a lesion is bypassed at the replication fork (such as the TT-CPD bypassed by Pol η), TLS is not in competition with DA, resulting in a predominance of TLS. In contrast, when the same lesion is bypassed post-replicatively, TLS competes with DA and the level of TLS is therefore reduced.

It is interesting to note that we obtained similar results with both constructs that express Pol η in G2/M. As stated earlier, *clb2-RAD30* afforded low-level expression of Pol η already in mid-to-late S phase. Given that our lesion is located close to an early replication origin (19), we expect it to be encountered by the replication fork early in S phase, when Pol η is still absent or at least very weakly expressed in the *clb2-RAD30* strain. Our observations suggest that TLS and DA compete, irrespective of whether Pol η reappears in mid-to-late S phase (*clb2-RAD30*) or in late S phase (*RAD30-AID\**), indicating that a daughter-strand gap has formed in both situations. This in turn suggests that if no TLS polymerase is capable of bypassing the lesion at the replication fork, repriming occurs quite rapidly, and bypass then occurs post-replicatively.

### TT(6-4), but not TT-CPD lesions lead to ssDNA accumulation during replication

If TT(6-4) lesions are bypassed post-replicatively, whereas TT-CPD lesions are bypassed at the fork, only the former should be accompanied by the generation of post-replicative gaps. To confirm this model, we generated strains that overexpress either yeast TT-CPD-specific photolyase (*ScPHR1* – noted “CPD Phr”) or *Xenopus laevis* TT(6-4)-specific photolyase (*xl64phr* – noted “6-4 Phr”) to selectively eliminate either TT-CPD or TT(6-4) lesions by photoreactivation. We confirmed that the photolyases were active in cells using dot-blots of DNA extracted from UV-irradiated cells overexpressing CPD Phr or 6-4 Phr with antibodies specific for the two UV lesions (Supp Fig 2). As a control, we used cells that express no photolyases (Phr-).

We exposed yeast cultures synchronized in the G1 phase to a single dose of UV (10 J/m^2^) (Figure 4A) and selectively eliminated either TT-CPD or TT(6-4) lesions by photoreactivation in the G1-arrested cells. Following release into S phase, we then monitored ssDNA by fluorescence microscopy using GFP-tagged Rfa1, the large subunit of the RPA complex (Figure 4C-D). We had previously shown that RPA foci serve as a proxy for post-replicative gaps (13). Cell-cycle stage was monitored by flow cytometry (FACS) (Figure 4B).

**Figure 4:**
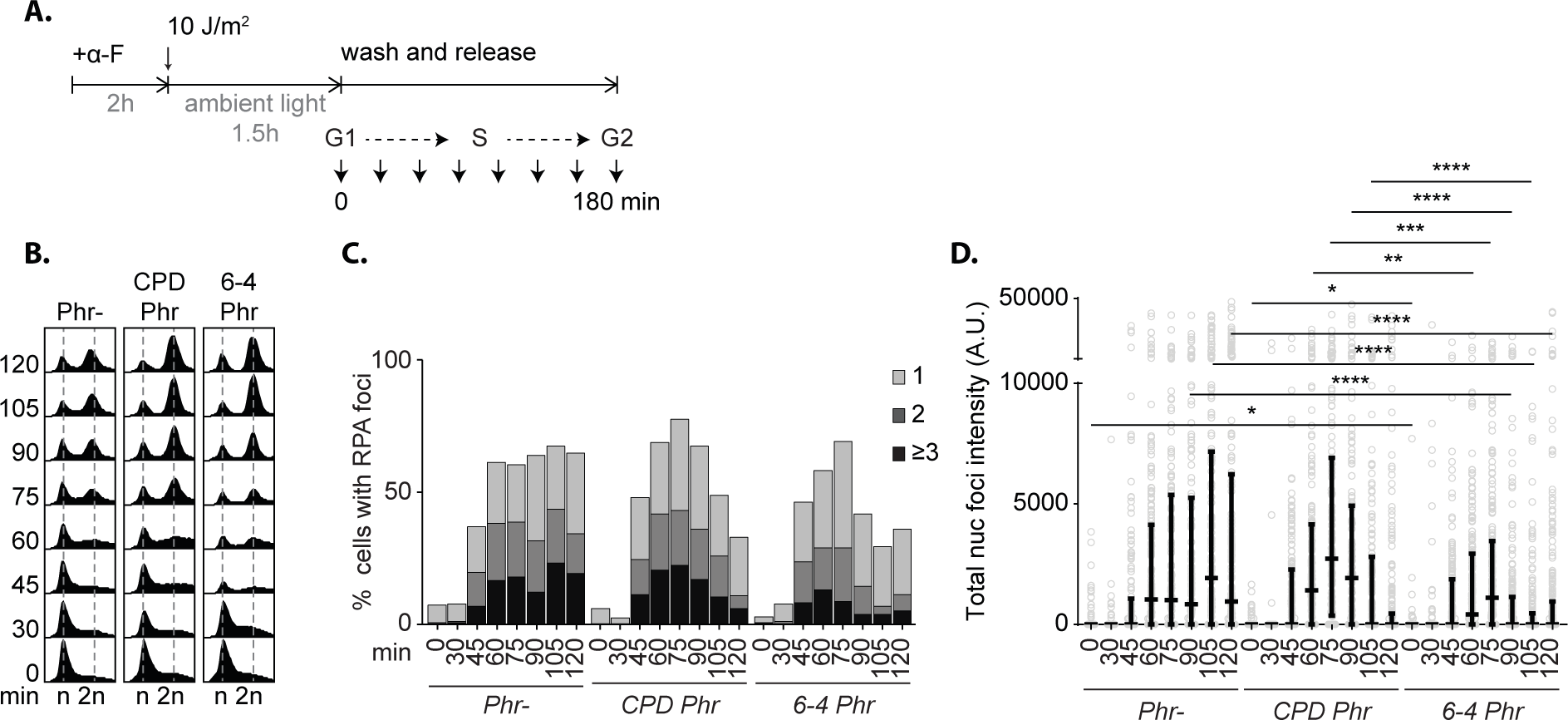
Experimental scheme: *rad14* cells arrested in G1 phase with alpha-factor (⍺F) were exposed to 10 J/m^2^ UV and photoreactivation by ambient light before release into the cell cycle. **B.** Cell cycle profiles monitoring progression through the cell cycle. **C.** Percentage of cells with Rfa1^GFP^ foci, detected by fluorescence microscopy. **D.** Total Rfa1^GFP^ foci intensity per nucleus. Mann-Whithney test ** p<0.01; ***p<0.001; ****p<0.0001.

In Phr-cells where both TT-CPD and TT-(6-4) lesions remained, and in cells in which TT-CPDs were repaired but TT(6-4) lesions remained (CPD Phr), we observed a higher number of foci arising per cell (Figure 4C) and a higher overall intensity of RPA foci per nucleus (Figure 4D), reflecting the total amount of ssDNA gaps generated at UV-induced lesions.

In contrast, in cells that had entered S phase with only TT-CPD lesions remaining (6-4 Phr), we observed fewer foci with an overall lower intensity per nucleus (Figure 4 C-D). It is important to note here that UV irradiation generates a much larger number of TT-CPD lesions compared to TT(6-4) lesions (26). Therefore, upon removal of TT(6-4) lesions, the total number of lesions remaining is likely much larger than upon removal of TT-CPDs. Nevertheless, despite the greater number of lesions remaining, we found the number and intensity of foci to be reduced. These results demonstrate that TT(6-4) lesions have a potent ability to induce post-replicative ssDNA gaps, while TT-CPD lesions do not.

### Gap extension by Exo1 promotes DA and reduces TLS

Our results suggested that the size of the post-replicative gaps could potentially play a crucial role in the choice of DNA damage tolerance pathway. To directly test this, we explored a possible contribution of *EXO1* to controlling the balance between DA and TLS. *EXO1* encodes a 5’->3’ exonuclease that has been mostly described for its role in recombination at double-strand breaks (during meiosis, but also in mitotic cells) and telomere maintenance (27). Additionally, Exo1 extends ssDNA gaps during Nucleotide Excision Repair (NER) (28). We have recently shown in bacteria that the extension of ssDNA gaps is crucial for DA to occur efficiently. Specifically, in the absence of the 5’->3’ exonuclease RecJ, we observed a decrease in DA and a concomitant increase in TLS (29, 30). The involvement of yeast *EXO1* in post-replication repair has been previously proposed (31-33), suggesting that gap extension is also required for DA in yeast. We also showed that Exo1 acts at post-replicative gaps to initiate damage signaling in response to MMS or UV treatment (34). We therefore set out to correlate the impact of *EXO1* on the size of the gaps with the choice of DNA damage tolerance pathway. By monitoring RPA foci, we first confirmed the contribution of Exo1 to gap widening in response to UV lesions in the absence of any photoreactivation (Figure 5). During early S phase, the number of cells with RPA foci and the number of foci per cell in response to UV irradiation did not differ between *EXO1* and *exo1* cells (Figure 5C). This is expected as these values correspond to the number of post-replicative gaps (directly correlated to the number of cells that have received replication-blocking damage) and thus should not be affected by the inactivation of *EXO1*. However, the overall intensity of RPA foci per nucleus, reflecting the total amount of ssDNA, was significantly lower in the *exo1* mutant (Figure 5D), indicating a reduction in the size of the post-replicative gaps. We therefore conclude that Exo1 extends the length of the ssDNA gaps generated at UV-induced lesions.

**Figure 5:**
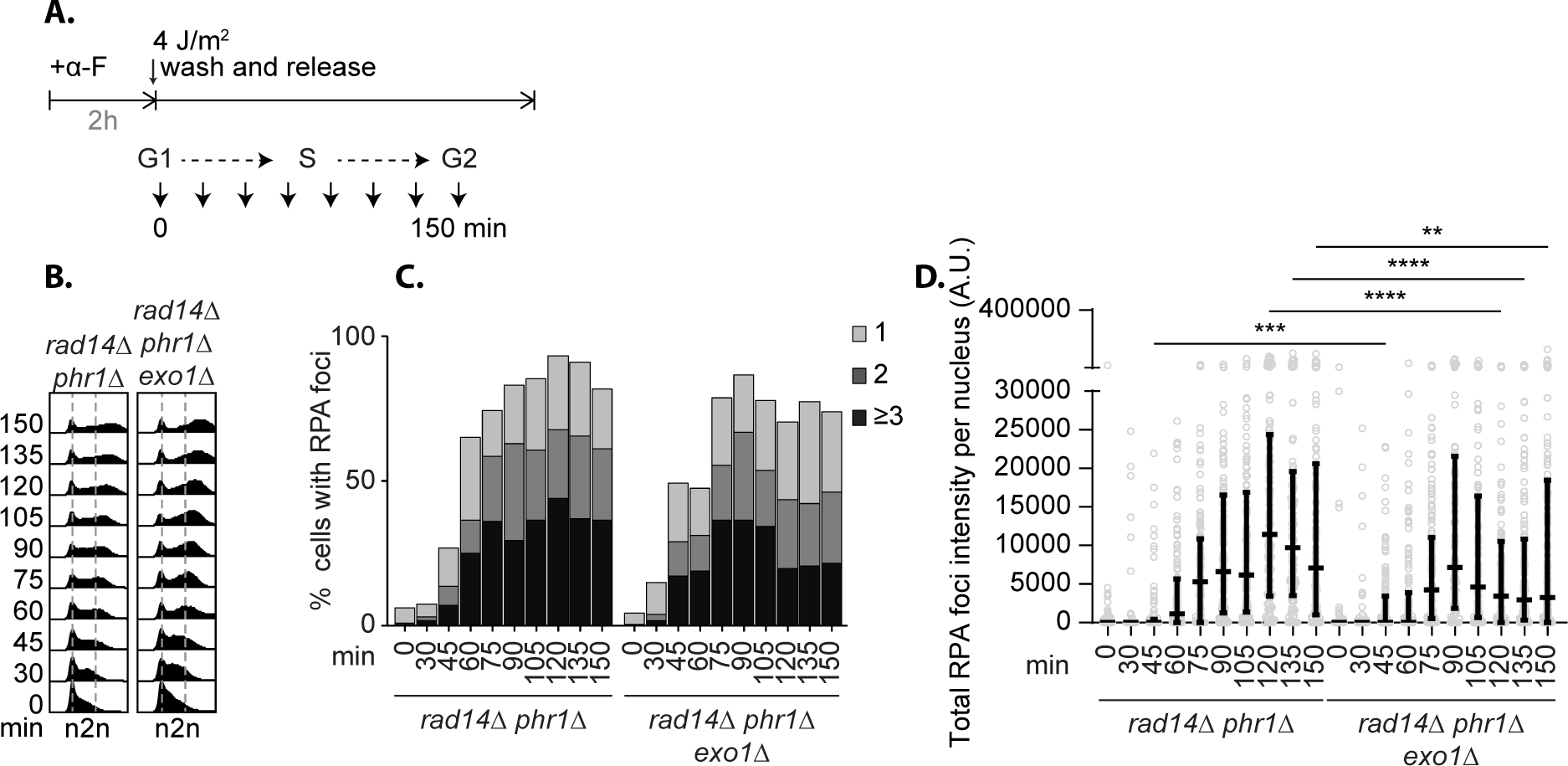
A. Experimental scheme: cells arrested in G1 phase with alpha-factor (⍺F) were exposed to 4J/m^2^ UV before release into the cell cycle without. **B.** FACS monitoring of the progression through the cell cycle. **C.** Percentage of cells with Rfa1^GFP^ foci, detected by fluorescence microscopy. **D.** Total Rfa1^GFP^ foci intensity per nucleus. Mann-Whitney test ** p<0.01; ***p<0.001; ****p<0.0001.

We then explored the effect of *EXO1* inactivation on the balance between TLS and DA at 3 different DNA lesions. As depicted in Figure 6, we observed a strong decrease in DA, accompanied with a strong increase in TLS for the TT(6-4) (>9 fold) and for the G-AAF (>3 fold) lesions in the absence of *EXO1*. Thus, in the absence of gap extension by Exo1, homologous recombination with the sister chromatid (DA) becomes less efficient, allowing TLS to occur at a higher rate. This confirms that TLS at these two lesions predominantly occurs at the post-replicative gaps.

**Figure 6:**
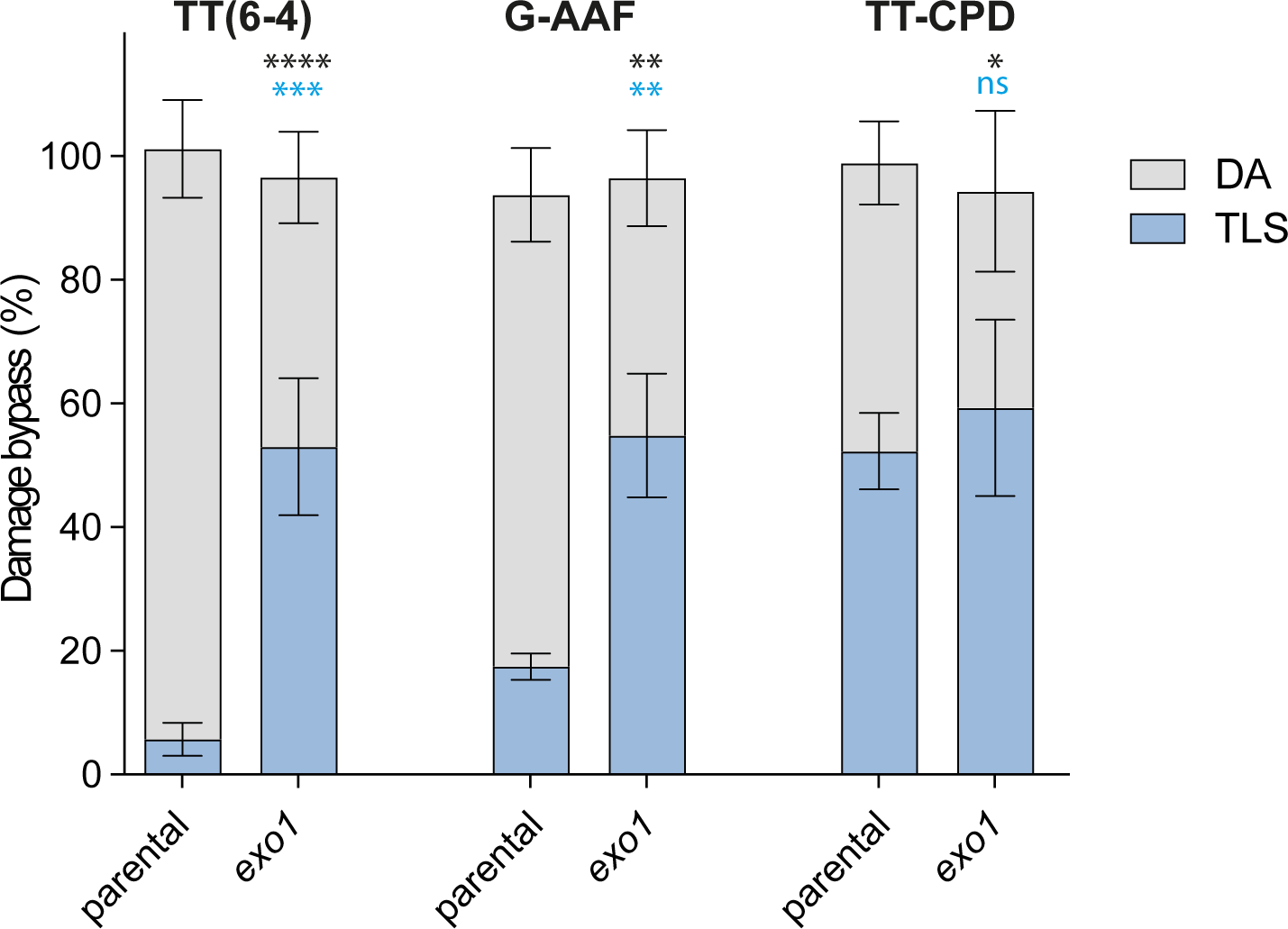
Partitioning of DNA damage bypass events at 3 different DNA lesion in the presence (parental) or absence of Exo1. Unpaired t-test was performed to compare TLS and DA values from the different mutants to the parental strain. (*p < 0.05; **p < 0.005; ***p < 0.0005).

In contrast, we did not observe any significant increase in TLS at the TT-CPD lesion in the absence of *EXO1*, confirming our model that gap extension has little effect on this lesion since it is mostly bypassed at the fork and not post-replicatively.

## Discussion

Our study reveals critical insights into DNA damage tolerance by examining the dynamics of translesion synthesis (TLS) and damage avoidance (DA) pathways at and behind the replication fork. Our findings highlight that TLS can occur both at the replication fork and at post-replicative gaps. We showed that DNA Pol η is able to bypass TT-CPD lesions directly at the fork, avoiding the generation of post-replicative gaps and thus the competition with DA. On the other hand, we showed that Pol ζ together with Rev1 bypass TT(6-4) and G-AAF lesions behind the fork, at post-replicative gaps. In this latter situation, TLS is in competition with DA and is therefore reduced. As a consequence, abolishing DA by the inactivation of *UBC13* leads to a strong increase in the usage of TLS (Figure 7).

**Figure 7:**
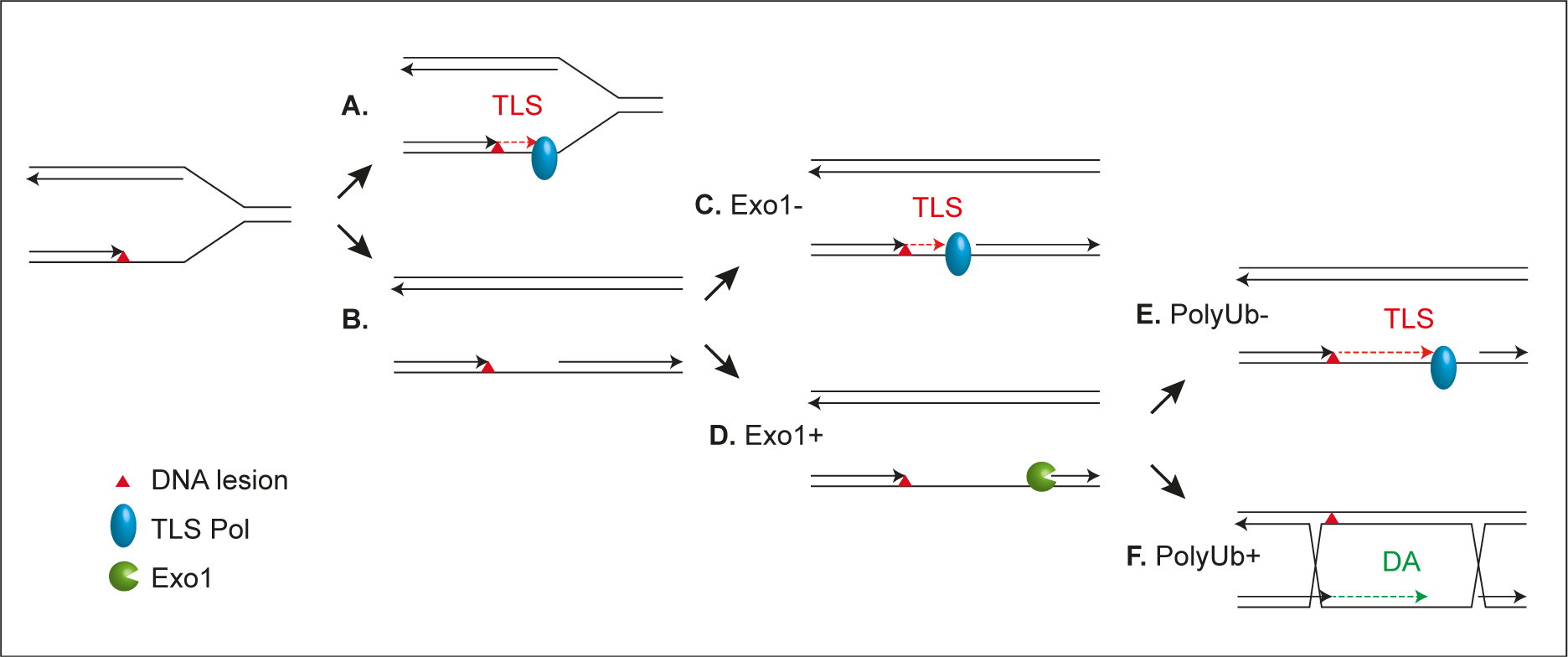
When the replication fork encounters a lesion, **A.** if a DNA polymerase able to bypass the lesion is present, TLS will occur at the fork. **B.** If no suitable polymerase is present, repriming will occur, generating a post-replicative gap. **C.** In the absence of Exo1, the gap will not be extended, preventing DA to occur and favoring TLS. **D.** When Exo1 is present, the post-replicative gap is extended. **E.** When PCNA cannot by poly-ubiquitylated (*ubc13*-), DA is inhibited, again leading to a strong increase in the use of TLS. **F.** Under conditions where PCNA is poly-ubiquitylated (*UBC13+*), DA will occur at the gap, therefore limiting the level of TLS.

The timing of expression of the TLS polymerases determines when TLS occurs. Rev1 is primarily expressed in G2/M (23, 24). Therefore, its activity, together with the activity of its interaction partner, Pol ζ (35), is limited to post-replicative gaps. This results in a reduced level of TLS by these two polymerases when DA is functional, as the two pathways are in competition. On the other hand, Pol η is expressed continuously throughout the cell cycle, allowing it to efficiently perform TLS at the fork during S phase without competing with DA. It would be interesting to test whether constant expression of Rev1 throughout the cell cycle would lead to an elevated level of TLS by enabling its action at the replication fork, thereby alleviating the competition with DA. However, the control of Rev1 appears not to be transcriptional (24), making its modulation challenging and such investigation difficult.

Using fluorescence microscopy, we showed that TT(6-4) lesions induced a higher number of post-replicative gaps as evidenced by the high number and intensity of RPA foci. This was not the case for TT-CPD lesions, indicating that these lesions are bypassed at the fork, avoiding the generation of post-replicative gap. These observations confirm that TT-CPD lesions are predominantly bypassed at the fork, whereas TT(6-4) lesions are bypassed at post-replicative gaps. This differential bypass of TT(6-4) and TT-CPD lesions appears to be conserved in human cells: Hung et al. have shown that TT(6-4) lesions are the key trigger for UV-induced ATR activation in mammalian cells, as these lesions induced ssDNA accumulation (36). Similarly, Benureau et al. have shown that Pol η prevents post-replicative gaps accumulation by bypassing TT-CPD lesions (37). Quinet at al. have shown that similarly, in mammalian cells, Pol ζ is responsible for the bypass of TT(6-4) at post-replicative gaps while Pol η bypasses TT-CPD lesion at the fork, as knockdown of Pol η enhanced fork stalling (38, 39). However, the role of Rev1 seems to differ in mammalian cells since the same study showed that Rev1 is required at the fork for TT-CPD bypass, but not at post-replicative gaps for TT(6-4) bypass (39). Similarly, in DT40 cells, Rev1 appears to be required at the fork and not at post-replicative gaps (15). These findings highlight a difference of functions for Rev1 between yeast and mammalian cells.

We also show that by promoting the extension of post-replicative gaps, Exo1 facilitates homologous recombination, favoring DA and in turn reducing TLS when the two pathways are in competition. It appears from our results, that for lesions such as TT(6-4) or G-AAF that are bypassed post-replicatively, the extension of the ssDNA gaps directly affects the balance between error-free and error-prone tolerance pathways. This reinforces the key role played by ssDNA gaps in genome instability, that has recently been pointed out by the Cantor group (40). The same group also showed how TLS plays an important role in filling these gaps (41).

From our observations, we can outline the following sequence of events in DNA damage tolerance (see Figure 7): During the S phase of the cell cycle, when the replication fork encounters a DNA lesion and stalls, TLS and repriming enter a competition. If a TLS polymerase able to bypass the lesion is available, TLS can occur at the fork (“on the fly”). This is the case for the TT-CPD lesion, which can be bypassed by Pol η as this polymerase is present is S phase. If Pol η is absent in S phase (or expressed later) or if the fork encounters a lesion such as TT(6-4) or G-AAF that cannot be bypassed by Pol η, TLS does not occur at the fork (Pol ζ-Rev1 being mostly active in G2/M). Instead, repriming will occur, generating a post-replicative gap. While the size of this gap remains to be determined, without extension by Exo1, the initial length of this gap is not enough to support efficient homologous recombination, and DA is inhibited, favoring TLS. Generally, when Exo1 is present, the gap is extended, supporting efficient DA while limiting TLS. At this latest stage, if DA is inhibited by abolishing PCNA poly-ubiquitylation (by the inactivation of *UBC13* for instance), TLS will again be favored.

Considering that TLS by Pol ζ-Rev1 at various lesions is much more mutagenic than TLS by Pol η at TT-CPD lesions (42), it is plausible that over evolution, cells might have restricted the expression of Rev1 to G2/M. This strategy might help to minimize mutagenesis by allowing competition with DA. On the other hand, direct, error-free bypass of the TT-CPD lesion by Pol η at the replication fork appears advantageous for the cell. TT-CPDs are the most abundant lesion directly induced by UV light, and their removal by NER is delayed compared to TT(6-4) (26). Allowing Pol η to act early in S phase thus prevents repriming and competition with other, less accurate TLS polymerases and again minimizes mutagenesis.

In conclusion, our findings show that cells potentially limit mutagenesis by permitting error-free TLS bypass directly at the fork, while restricting error-prone bypass to post-replicative gaps where it competes with error-free DA. This dual approach allows for efficient damage tolerance while minimizing the risk of mutagenic events.

## Materials and methods

### Yeast strains

All strains were cultured in YPD or low fluorescence synthetic complete (SC) media supplemented with appropriate amino acids at 30°C. All strains used in the present study are derivative of strain EMY74.7 (43) (MATa *his3-Δ1 leu2-3,112 trp1Δ ura3-Δ met25-Δ phr1-Δ rad14-Δ msh2Δ:hisG*). In order to study tolerance events, all strains are deficient in repair mechanisms: nucleotide excision repair (*rad14*), photolyase (*phr1*), and mismatch repair system (*msh2*). Gene disruptions were achieved using PCR-mediated seamless gene deletion (44) or URAblaster (45) techniques. Promoter replacement and fluorescent tags were introduced with standard PCR-based methods (46).

Strains used in photoreactivation experiments were overexpressing yeast *PHR1*, encoding TT-CPD photolyase (pGPD promoter replacement in the native locus), or *Xenopus laevis* x64lphr 64PP photolyase (integrative plasmid pHU5768 carrying pADH_xl64phr integrated into the *LEU2* locus). A plasmid carrying yeast codon-optimized xl64phr used for cloning was synthesized by Twist Bioscience.

Strains carrying *RAD30* (encoding Pol η) under control of an auxin-inducible degron (AID*) tag were created by inserting the pKAN-PRAD30-9myc-AID*(N) cassette into the *RAD30* native locus.

Strains carrying *RAD30* under control of regulatory elements of cyclins Clb2 (from pGIK43) or Clb5 (from pKM101) were created by inserting a cyclin cassette into the native *RAD30* locus. TIR1 strains were created by integration of the osTIR1 cassette from pNHK53 (encoding OsTIR1 under control of the *ADH1* promoter) into yeast chromosome VII.

Plasmid pNHK53 was obtained from the National BioResource Project–Yeast (47), and plasmid pGIK43 from Georgios Karras. Plasmid pKAN-PCUP1-9myc-AID*(N) was previously described (25).

### Integration

Integration of plasmids carrying TT(6-4) or G-AAF lesions (or control plasmids without lesion) was performed as previously described (19).

For experiments involving cell-cycle restricted Pol η, cells were synchronized in the G1 phase using alpha-factor. After synchronization, the washing, conditioning and electroporation steps were carried out directly. For strains with AID* degrons, auxin was present at 1 mM during incubation with alpha-factor.

Lesion tolerance rates were calculated as the relative integration efficiencies of damaged vs. non-damaged vectors, normalized by the transformation efficiency of a control plasmid (pRS413) in the same experiment. DA events are calculated by subtracting TLS events from the total lesion tolerance events. All experiments were performed at least in triplicate. Graphs and statistical analysis were performed using GraphPad Prism, applying unpaired t-tests. Bars represent the mean value ± s.d. (standard deviation).

### Detection of proteins

Total lysates of synchronized yeast cultures were prepared by quick trichloroacetic acid (TCA) extraction: cells (pelleted 10 ml of culture) were resuspended in 250 μl of 20% TCA and vortexed with glass beads for 30s. After centrifugation at 3000 × g for 10 min, the supernatant was removed and the pellet resuspended in LDS loading buffer and incubated at 75°C for 10 min. Proteins were analysed by SDS–PAGE/Western blotting using monoclonal antibodies against c-Myc (9E10) or HA (12CA5, Thermo).

### Cell cycle analysis

Cells were fixed in 70% ethanol overnight and washed twice with 50 mM TE, pH 7.5. After incubation with 0.1 mg/ml DNase-free RNAse A for 4 h at 42°C, and 0.5 mg/ml Proteinase K for 30 min at 50°C, samples were sonicated on low setting for 10x 3 s and DNA was stained using 1 µM SYTOX green. DNA content was analyzed by flow cytometry using a Accuri C6 Plus flow cytometer (BD Biosciences).

Alternatively, cells were fixed in 70% ethanol and washed twice with 50 mM sodium citrate pH 7.0. After incubation with 80 mg/ml RNase A at 50°C for 1 h, and with 80 mg/ml Proteinase K at 50°C for 1 h, samples were stained with 32 mg/ml propidium iodide and sonicated on a low setting for 3 s. DNA content was analyzed by flow cytometry with a Novocyte Quanteon (Agilent).

### Live cell imaging and image analysis

Imaging and image analysis were performed in duplicate, essentially as previously described (13). Cells synchronized in G1 with alpha-factor were irradiated with UV, washed, and released into the cell cycle. For experiments involving photoreactivation, cultures were incubated immediately after UV irradiation with agitation in open Petri dishes placed 5 cm below a lamp (daylight LED lamp, LeuchtenDirekt, 24 W, 5000K) for 90 min at room temperature, with alpha-factor.

At indicated time points after release, cells were plated on concanavalin A-coated chambered coverslips with glass bottom (Ibidi) and imaged with a DeltaVision Elite system (GE Healthcare) equipped with a 60× oil immersion objective (NA = 1.42), scientific CMOS camera, InsightSSI solid state illumination, and SoftWoRx software with built-in deconvolution algorithms in an environmentally controlled chamber at 30°C. GFP signals were imaged with a FITC filter and DIC was used for brightfield images. Z stacks with 21 planes (step size = 0.2 μm) were acquired for each image.

Images were analyzed using customized scripts (13) written in ImageJ macro language with ImageJ FIJI software (https://fiji.sc/). Scripts are available at https://github.com/helle-ulrich-lab/image-analysis-PORTs.

### Detection of TT-CPD and 64-photoproduct lesions by dot blot

10 ml of UV-treated yeast cultures OD_600_=1 (with or without photoreactivation) were harvested. Genomic DNA was isolated using MasterPure™ Yeast DNA Purification Kit (Biozym) according to the manufacturer’s protocol. An appropriate amount of heat-denaturated DNA (1.5 µg for TT-CPD detection and 6 µg for 6-4PP detection) was spotted onto HybondN+ membrane (Cytiva). Samples were fixed by baking (80°C, 2 h). Lesions were detected using antibodies: anti-CPD (Cyclobutane Pyrimidine Dimer) (clone TDM-2; #NM-DND-001; Cosmo), or anti-6-4 PPs (6-4 photoproducts) (clone 64M-2; #NM-DND-002; Cosmo).

## Acknowledgements

We thank Luisa Laureti for critical reading of the manuscript.

