## Supplementary material for "Post-replicative lesion processing limits DNA damage-induced mutagenesis": suppl figure 1 and 2

### Cells limit mutagenesis by processing DNA lesions behind the replication fork

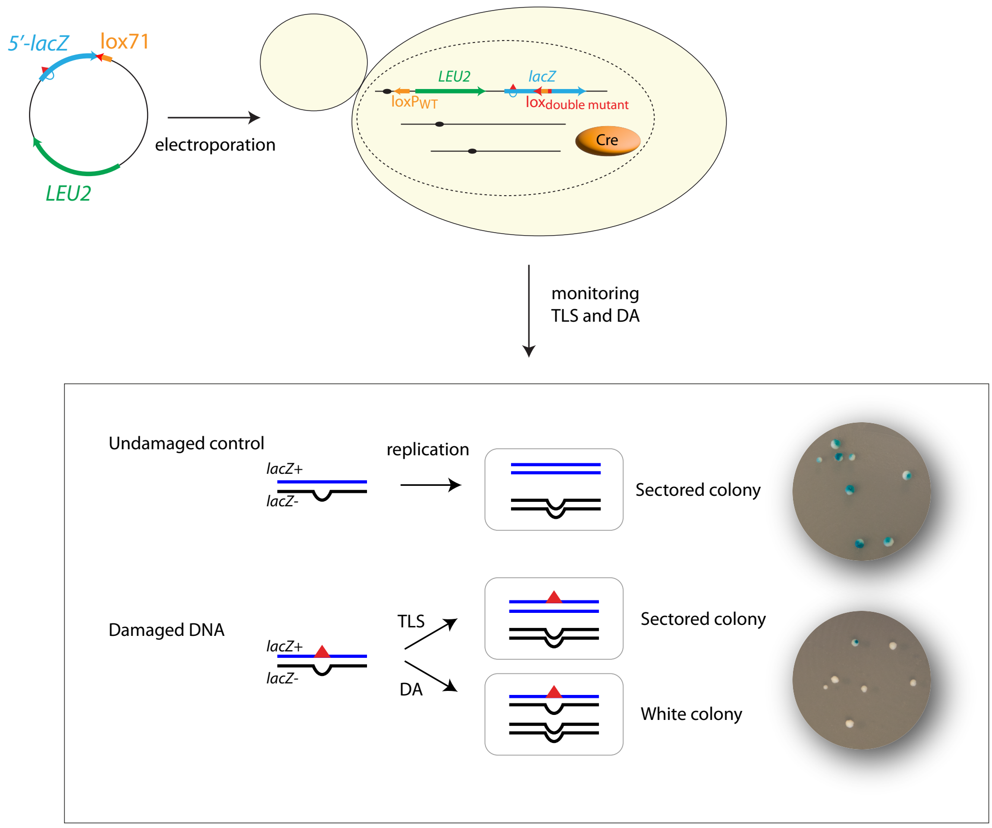


**Supplementary Figure 1:** outline of the integration system: a non-replicative plasmid containing a single lesion is integrated into one of the yeast chromosomes using Cre/lox site-specific recombination. The integrative vector carrying a selection marker (*LEU2*) and the 5′-end of the *lacZ* reporter gene containing a single lesion is introduced into a specific locus of the chromosome with the 3′-end of *lacZ*. The precise integration of the plasmid DNA into the chromosome restores a functional *lacZ* gene, enabling the phenotypical detection of TLS and DA events (as blue and white colonies on X-gal indicator media).

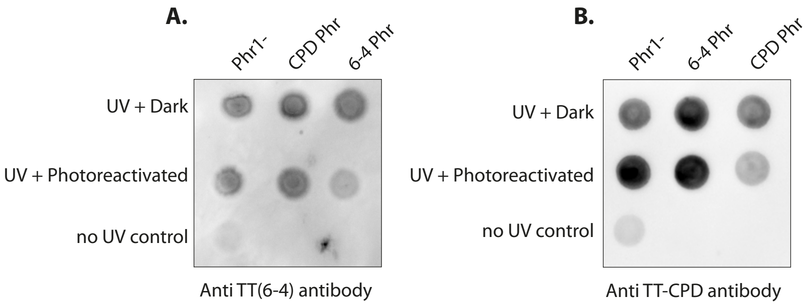


**Supplementary Figure 2:** Dot-blot validation of lesion-specific repair by photoreactivation. Cells expressing no photolyases (Phr-), yeast TT-CPD-specific photolyase (*ScPHR1* – noted "CPD Phr"), or *Xenopus laevis* TT(6-4)-specific photolyase (*xl64phr* – noted "6-4 Phr") were UV irradiated (10J/$m^2$), and either photoreactivated or kept in the dark. Genomic DNA was extracted, quantified, heat denatured, and then spotted onto membranes. Membranes were probed with **A.** anti-TT(6-4) antibodies or **B.** anti-CPD antibodies. *PHR1* and *xl64phr* photolyases efficiently repair their cognate lesions.
